# Deep-Learning-Based Multivariate Pattern Analysis (dMVPA): A Tutorial and a Toolbox

**DOI:** 10.1101/2020.12.03.410910

**Authors:** Karl M. Kuntzelman, Jacob M. Williams, Phui Cheng Lim, Ashok Samal, Prahalada K. Rao, Matthew R. Johnson

## Abstract

In recent years, multivariate pattern analysis (MVPA) has been hugely beneficial for cognitive neuroscience by making new experiment designs possible and by increasing the inferential power of functional magnetic resonance imaging (fMRI), electroencephalography (EEG), and other neuroimaging methodologies. In a similar time frame, “deep learning” (a term for the use of artificial neural networks with convolutional, recurrent, or similarly sophisticated architectures) has produced a parallel revolution in the field of machine learning and has been employed across a wide variety of applications. Traditional MVPA also uses a form of machine learning, but most commonly with much simpler techniques based on linear calculations; a number of studies have applied deep learning techniques to neuroimaging data, but we believe that those have barely scratched the surface of the potential deep learning holds for the field. In this paper, we provide a brief introduction to deep learning for those new to the technique, explore the logistical pros and cons of using deep learning to analyze neuroimaging data – which we term “deep MVPA,” or dMVPA – and introduce a new software toolbox (the “Deep Learning In Neuroimaging: Exploration, Analysis, Tools, and Education” package, DeLINEATE for short) intended to facilitate dMVPA for neuroscientists (and indeed, scientists more broadly) everywhere.

## 1 Introduction

Although the roots of cognitive neuroscience date to the 1920s (the advent of electroencephalography, EEG; Berger, 1929), the modern neuroimaging era began in the mid-1990s, with the development of functional magnetic resonance imaging (fMRI) methodology and the increasingly widespread availability of (affordable) desktop computing workstations powerful enough to process fMRI datasets. In those days, data analysis was primarily limited to univariate investigations such as event-related potentials (ERPs) in EEG and univariate general linear model (GLM) analyses aimed at detecting “blobs” of activation with fMRI (as well as differences in activity, e.g. between experimental conditions, within such blobs)^1^. However, the march of progress towards ever-more sophisticated models of brain function and the testing of ever-more refined hypotheses has created a demand for corresponding improvements in analysis techniques.

Thus, somewhat more recently (beginning in the early-to-mid-2000s), a second age in neuroimaging analysis arose with the advent of multivariate pattern analysis (MVPA; Haxby et al., 2001; Haxby et al., 2014). Rather than focusing on whether a certain cognitive event elicits activity in a particular cluster of fMRI voxels (or a voltage peak at a particular temporal latency with ERP), MVPA is instead concerned with how a neural pattern or multivariate “brain state” comprising multiple voxels (fMRI) or electrode/timepoint combinations (EEG) might collectively correspond to a certain cognitive event or state. Numerous MVPA variations exist, including those based on correlation (either Pearson or rank-based; Haxby et al., 2001), support vector machines (SVMs; De Martino et al., 2008; Dosenbach et al., 2010), logistic regression (Akama et al., 2012), sparse multinomial logistic regression (SMLR; Kohler et al., 2013; Krishnapuram et al., 2005), naïve Bayes classifiers (Kassam et al., 2013), and more. Many of these techniques concern classification of brain patterns into discrete cognitive states, whereas others examine different aspects of the data (e.g., overall similarity between brain patterns; Xue et al., 2010; Lim et al., 2019) without explicit categorization, but all of them represent increases in mathematical and conceptual sophistication over univariate techniques. Importantly, when compared to earlier univariate techniques, MVPA has enabled us to examine in a much more nuanced fashion how brain activity patterns encode mental states.

Although traditional MVPA techniques are substantially more advanced than univariate techniques, they are nonetheless still fairly simple, both mathematically and conceptually. Traditional MVPA is a form of machine learning (ML), but it is among the simplest forms; most MVPA approaches use straightforward linear mathematical models. This comparative simplicity certainly confers advantages – for example, faster computation times than more complex techniques (with some caveats^2^), and a generally lower risk of “overfitting”^3^. However, simpler mathematical formulations are necessarily limited in what we call “informational resolution” – the specificity of the neural patterns and cognitive states that they are able to capture.

How much informational resolution is required to glean as much about brain function as is possible using current neuroimaging technology? The answer is hard to pin down, partly because it is difficult to establish firm estimates of the “noise ceiling”^4^ for these techniques. As neuroimagers, we often complain that our techniques are “noisy,” but with proper usage, the signal-to-noise ratios of EEG and fMRI are really rather high, when considering only measurement noise from the instruments themselves and the surrounding physical environment. Of significantly greater concern are “noise” sources such as subject head/body motion, physiological artifacts (cardiac, respiratory, muscular, etc.), and cognitive artifacts (distraction, poor understanding of instructions, falling asleep). Noise ceilings for certain analytic techniques and datasets can be estimated (Kay et al., 2008; Nili et al., 2014), but ultimately they will depend on which data components are considered “noise”; aside from the noise that arises from the physics of the measurement itself, other biological and subject-driven artifacts have some hope of being detected, modeled, and/or removed. And, much like the signal components we actually care about (i.e., those related to our experimental questions), our ability to detect and account for noise depends largely on the sophistication of our analytic techniques.

What we do know is that the brain is a highly complex, highly nonlinear system (Koch & Laurent, 1999; Sporns et al., 2000; Buzsaki & Mizuseki, 2014), and the addition of noise sources that are also complex and nonlinear makes brain data no easier to analyze and interpret. Although the limits of the usefulness of traditional MVPA, with its relatively low informational resolution, have not yet been reached, those limits do loom on the horizon. As the size of neuroscience data continues to grow^5^, traditional MVPA’s limitations become ever more apparent. It is a statistical truism that more complex analytic models, with more parameters to fit, allow us to account for a greater proportion of a dataset’s variance, but they also require larger input data to estimate their parameters reliably. Yet the sizes of many contemporary datasets are now such that they can potentially accommodate significantly more sophisticated statistical models than traditional MVPA, with greater power to identify, extract, and distinguish noise sources and signals of interest. Thus, we believe it is time for cognitive neuroscience and related fields to place increased emphasis on developing, exploring, and using more sophisticated techniques, and on producing tools that can be used to perform that exploration more effectively and efficiently.

### 1.1 The case for deep learning

There are numerous potential analytic methods of greater complexity and sophistication than traditional MVPA. One class of ML techniques that has been gaining popularity, and the one we endorse in this paper, is “deep learning.” Deep learning, briefly defined, refers to the use of artificial neural networks (ANNs), typically with recurrent and/or convolutional architectures, that are more complex, flexible, and powerful than both earlier generations of ANN architectures and the techniques used for traditional MVPA. In the last few years, such deep neural networks (DNNs) have been used increasingly heavily in a number of fields that employ ML for all kinds of purposes. Such usage includes an ever-growing collection of studies in human neuroscience and related disciplines, although a relatively small proportion have been devoted to neuroimaging analysis, and fewer still devoted to decoding cognitive states from functional measurements of brain activity, which is a topic of great interest to many. We believe the studies so far represent only the tip of the proverbial iceberg in terms of what is achievable by using DNNs to analyze neuroscience datasets. In fact, we believe deep learning has the potential to perform most of the tasks for which traditional MVPA is typically employed, but with greater speed, flexibility, and power, and thus we advocate for the more widespread use of what we call “deep MVPA,” or dMVPA for short.

To achieve more widespread adoption of deep learning in the neurosciences, notable challenges to confront include 1) a relatively low level of knowledge/awareness of these techniques, and 2) insufficient availability of software tools to make dMVPA as approachable as traditional MVPA. In this paper we address the first challenge by providing a brief review of deep learning techniques, including how they can be used in neuroscience investigations, and the pros and cons of dMVPA versus traditional MVPA. We address the second challenge by introducing a new Python-based software toolbox (the “Deep Learning In Neuroimaging: Exploration, Analysis, Tools, and Education” package; DeLINEATE for short) that builds upon previous DNN and MVPA tools and aims to make dMVPA more approachable and efficient for other researchers.

## 2 dMVPA: A tutorial

### 2.1 A brief history of neural networks

The techniques we now collectively call “deep learning” are generally extensions of older “shallow” ANNs, which are significantly less complex and powerful than DNNs but not much different in their basic principles. The concept behind all ANNs originates from a highly abstracted view of non-artificial neural networks, i.e., the biological nervous system (Figure 1A). In this framework, most implementation details are stripped away, and what remains is the basic idea of a network of simple computational units (“neurons”) that receive input (which can typically be excitatory or inhibitory), perform an operation on their inputs (typically some variation on summation), and produce an output (typically a single value analogous to an action potential or a firing rate), which might then serve as input to one or more downstream neurons^6^. The original and simplest case is the McCulloch-Pitts neuron (McCulloch & Pitts, 1943; Figure 1B), a processing unit whose input and output values are exclusively binary (0 or 1). The McCulloch-Pitts neuron sums its inputs, compares the sum to some threshold value, and outputs a 1 (“action potential”) or 0 according to whether the sum exceeds the threshold. Although a pioneering idea and an interesting (if highly simplified) early model of neural information processing, McCulloch-Pitts neurons can only implement a limited set of functions and are thus not considered very useful for modern ML applications.

**Figure 1.**
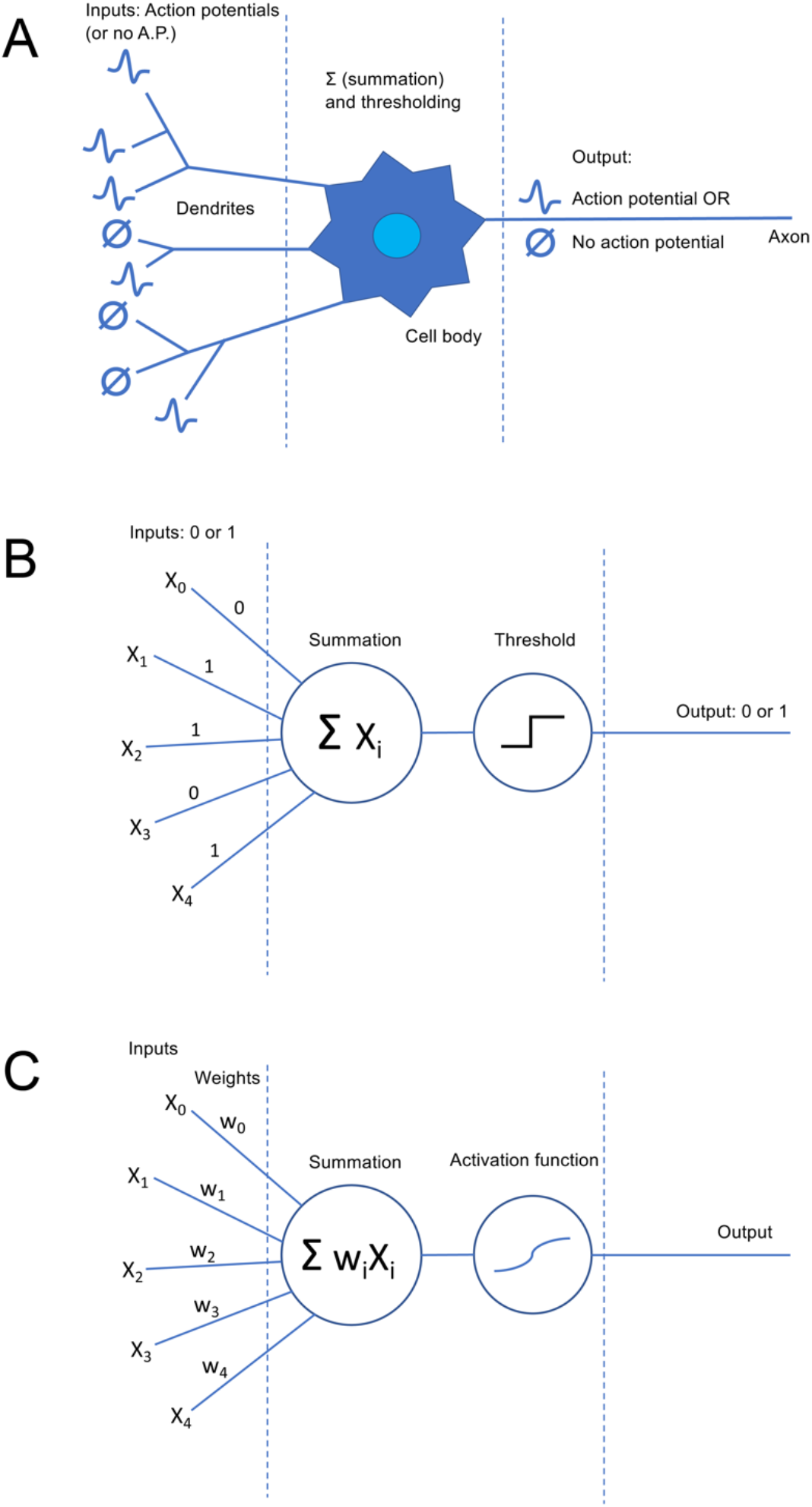
Comparison of biological and artificial neural models. (A) A simplified “textbook” model of a biological neuron. Inputs come in via the dendrites in the form of action potentials (or the lack thereof). The inputs are summed in the cell body (soma) and, if the threshold voltage is reached, the cell produces an action potential as output that is delivered via the cell’s axon. (B) The original and simplest version of an artificial neuron model, the McCulloch-Pitts neuron. Similar to the biological neuron, inputs (xi) and outputs are binary (although we now know this to be an oversimplified view of biological neurons). Inputs are summed and the result passed to a thresholding function; if the threshold is met, an output of 1 is produced, and otherwise the output is 0. (C) A perceptron, a more sophisticated revision of the McCulloch-Pitts neuron that has an important place in modern artificial neural networks. The concept of trainable weights (wi; mimicking biological potentiation at synapses) is introduced, and inputs are now multiplied by their corresponding weight before summation. In addition, in contemporary perceptron models, the threshold function can be replaced by any arbitrary function, called the “activation function.” Popular activation functions like the hyperbolic tangent may still act largely like thresholding functions, but with the ability to deliver graded rather than strictly binary output values.

A few years later, though, ANNs took a significant step forward when Rosenblatt (1958) incorporated Hebb’s theoretical views on the strengthening and weakening of synaptic connections (Hebb, 1949) into a McCulloch-Pitts-like unit that came to be called the *perceptron*^7^. In its simplest form, a perceptron (Figure 1C) is largely identical to a McCulloch-Pitts neuron with one critical addition: Each input is now associated with a “synaptic weight” (often denoted *w_0_, w_j_*, etc.) that determines whether it is excitatory or inhibitory and how strongly it influences the output. Summation is then performed on the inputs after they have been multiplied by their respective weights. Statistically inclined readers may recognize this as not-dissimilar-to a regression model, particularly logistic regression; to conceptually convert between a multiple regression model and a perceptron, simply rename the weights from the *β_i_* typically used in regression equations to *w_i_* and pass the regression output through a thresholding function, logistic function (to essentially replicate logistic regression), or other function as desired^8^. This function is known as the artificial neuron’s *activation function;* activation functions are a key feature of contemporary ANN designs, and there are many options to choose from.

In most important ways, the perceptron-like artificial neural units used in some DNNs today are not substantially different than the classical perceptrons discussed by Minsky and Papert (1969) in their seminal book some 50 years ago. Yet the original perceptron architectures retained many of the McCulloch-Pitts neurons’ limitations and still had significant constraints on the classes of problems they could solve. The key developments that distinguish the powerful deep learning techniques of today from the toy models of the past are 1) improved methods for establishing what the proper synaptic weights should be for a given dataset/problem, i.e., *training* the neural network, and 2) new and ultimately better ways of digitally connecting groups of artificial neurons together into more complex structures, i.e., improved ANN *architectures*^9^.

### 2.2 Training algorithms and neural network architectures

The earliest ANN architectures were very simple indeed; either a single artificial neuron or, in the next major architectural advance after that, a *layer* of such units. In this latter (still very simple) architecture, the units are *fully connected*, meaning that each unit receives a copy of each possible input value (see Figure 2A). Note that in this figure, as in many neural network diagrams, inputs and outputs are represented as “layers” of a sort, but there is only one true layer of computational units^10^. If the ANN is meant to calculate a classification problem (a common application), the outputs are typically assumed to each correspond to one of the possible classes, and are interpreted in a winner-take-all fashion (i.e., for a given set of input data, whichever output value is highest is interpreted as the network’s prediction of the class that the input data belong to). Although the transition from single-neuron to single-layer architectures laid a critical foundation for later work, single layers of perceptrons were soon shown not to be terribly useful as artificial intelligence agents, no matter what their synaptic weights were or how those weights were determined. As Minsky and Papert demonstrated in *Perceptrons* (1969), it is mathematically impossible for any single-layer perceptron network – no matter how many units are in it – to perform certain fundamental computational operations^11^.

**Figure 2.**
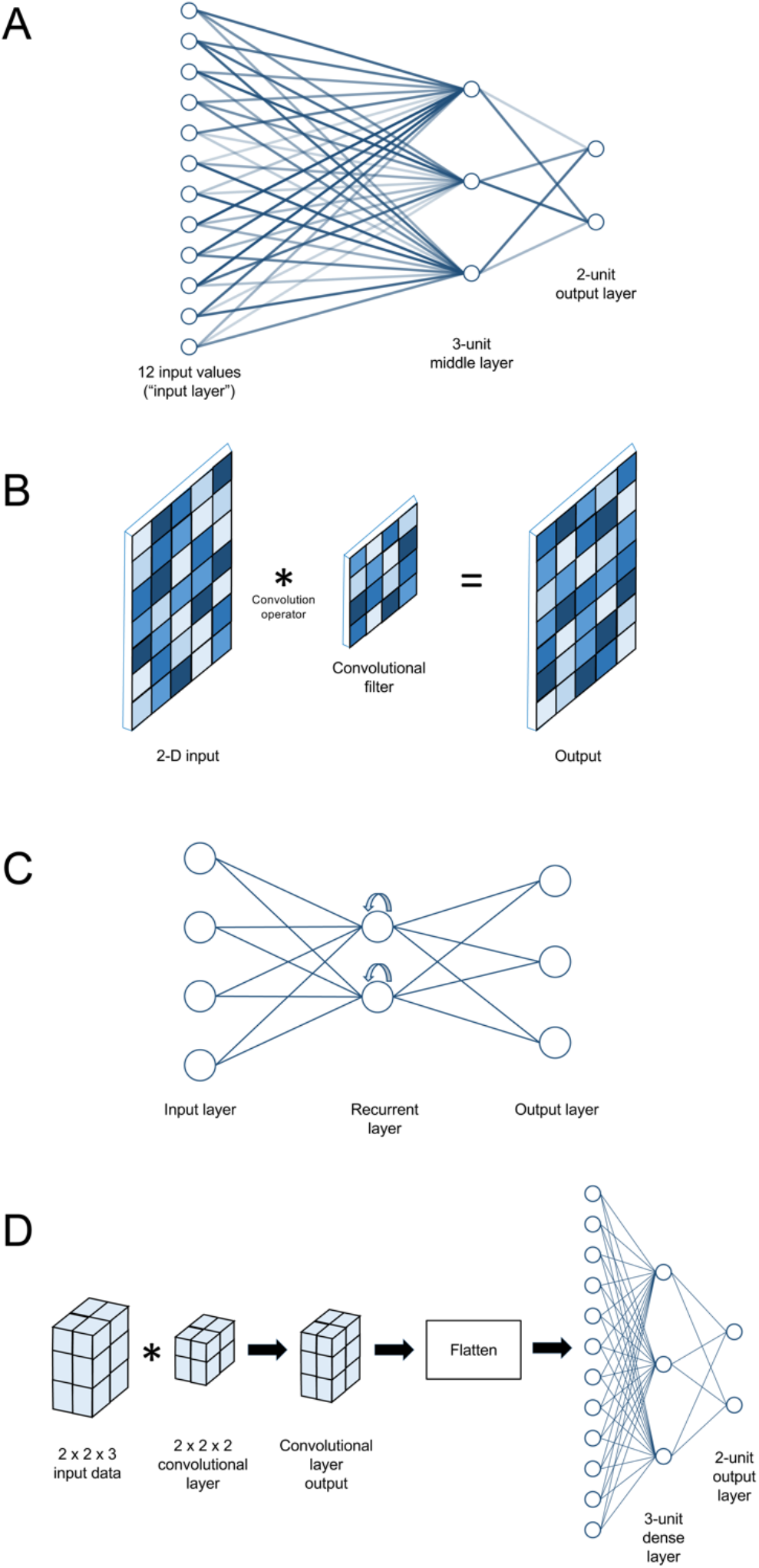
Examples of artificial neural network architectures. (A) A simple fully-connected multi-layer perceptron model with 12 input values, a middle layer comprising three perceptrons, and an output layer with two perceptrons. Line lightness is used to represent synaptic weight strength. (B) An example of a convolutional neural network layer that might be used to analyze 2-D input. Here, the layer looks less like a set of artificial neurons and more like a digital filter used in image processing. Two-dimensional input is convolved with a 2-D filter to yield a 2-D output, sized similarly to the input. During training, it is the values in the convolutional filter that get adjusted. Square lightness represents the numeric values in the cells of each 2-D matrix. (C) An example of a simple recurrent neural network. There are many types of recurrent neural network structures with varying degrees of complexity, but all share the property that recurrent units’ output gets passed back into them (represented here by curved arrows), giving them some form of “memory” for previous input values. (D) An example of a complete neural network architecture that might be used to analyze 3-D input such as MRI data for a two-class classification problem. In this simple example, 12 input values in a 2×2×3 array are first passed through a 2×2×2 convolutional filter, yielding another 2×2×3 array as output. This is then passed through a “flattening” layer to convert it to a 12×1 vector, which then passes through a 3-unit dense layer to a 2-unit output layer (as shown in panel A).

This revelation may not seem surprising in retrospect; after all, a single layer of neurons, all receiving the same inputs, is not a very viable architecture for a biological neural network either. Still, it was enough to significantly dampen enthusiasm for ANN research for over a decade. Although adding another layer of computational units (known as a *hidden layer*) would allow the network to maintain an intermediate representation of the input and enable more complex operations^12^, the algorithms available for training single-layer perceptron networks could not be readily extended to multi-layer architectures. In the 1980s, however, interest was reignited with the popular (re-)discovery of the *backpropagation* algorithm (or simply *backprop*, to its friends). This algorithm was known and even applied to ANNs previously (Linnainmaa, 1970; Werbos, 1974), but it did not reach mainstream awareness until the publication of Rumelhart and colleagues’ (Rumelhart et al., 1986a; Rumelhart et al., 1986b) seminal formulations of it. Backprop proved to be a highly robust method for training ANNs across many applications, and is still the dominant training algorithm in use today.

The main principle behind backprop is to take any errors made by the network during training and propagate responsibility for them from the output layer (where the error is assessed, by comparing the network’s decision to the known correct decision^13^) backwards through the network towards the input layer, penalizing the synaptic weights most responsible for the error along the way. It is analogous to the human behavior encapsulated by the vernacular phrase, “Shit rolls downhill.” For example, imagine that a CEO – the final decision-maker in her company’s chain of command – makes a decision that loses the company money. She turns to her immediate inferiors and doles out punishment to them proportional to how influential they were in guiding that decision, and decides to trust those influential individuals less in the future. In turn, each of those upper-level managers passes along the punishment and distrust they have received to their immediate inferiors, again proportional to their influence on the upper managers’ actions, and so on down the corporate hierarchy. In this way, one hopes that the next time a similar decision is faced, the shift in influences and communication channels throughout the hierarchy will produce a better outcome.

The advent of effective backprop-based training for ANNs reignited interest in them for a time, and backprop-trained ANNs were found to perform admirably in a number of ML domains. Still, before long, interest waned again, as neural nets with many hidden layers were found to present mathematical difficulties for backpropagation algorithms, and complex networks also took a long time to train on the CPUs of the era. Concurrently, the 1990s also saw the development of promising alternative ML algorithms, most notably the modern incarnation of support vector machines (SVMs; Cortes & Vapnik, 1995; Boser et al., 1992). SVMs were easier to work with than ANNs and performed nearly equivalently (or even better) in many problem domains of the time. Thus, when traditional MVPA techniques arose in neuroimaging in the 2000s, it is unsurprising that SVMs and other similarly robust linear classification algorithms, well-suited to the mid-sized datasets of the time, dominated within that emerging field.

### 2.3 The deep learning Renaissance

Research interest in ANNs experienced another upswing, which has continued to the present, beginning around 2006. This rebirth happened for several reasons, including: 1) solutions to some of the technical and mathematical problems that had plagued networks with complex, many-layered architectures (Hinton et al., 2006); 2) methods for training ANNs on desktop workstations using the GPU instead of the CPU, producing speed improvements of up to ~70x (Raina et al., 2009); 3) the advent of the so-called “Big Data” era, which provided the larger datasets required to adequately train more complex neural architectures; and 4) the re-branding of neural net research as “Deep Learning,” which, despite being more public relations than true substance, still likely helped ignite new interest in a field formerly seen as relatively tired and unpopular. Since this Renaissance began, there have naturally been several key architectural and methodological developments^14^. However, these newer architectures are still trained and used similarly to the older, simpler networks described above, and the variations are not too difficult to comprehend once one understands the fundamental concepts and terminology behind ANNs.

During the early days of this revival, deep learning research had a number of notable successes, including advances in speech recognition, natural language processing, computer vision, financial fraud detection, and more. Large technology companies, who had access to Big Data and financial motivations for finding better ways to process it, also had their interest piqued. Thus they began to invest in deep learning research themselves, including developing improved software tools (for example, the TensorFlow toolbox, developed primarily at Google, and PyTorch, developed primarily at Facebook). These tools typically rely on lower-level driver and software library support for GPU-based computation, most notably NVIDIA’s CUDA libraries for general GPU-accelerated computing and their cuDNN framework, built atop CUDA, specifically for DNN applications^15^. Although the use of such tools has exploded in the technology sphere and in basic computer science research, adoption in other areas, such as cognitive neuroscience, has been slower. This lag can partly be attributed to fundamental limitations and difficulties of DNN-based data analysis (e.g., potential for overfitting), but another large factor is the lack of higher-level software tools that make it convenient for neuroscience researchers to implement dMVPA without needing to write large amounts of their own code. And, when better software tools exist, it will be more efficient to explore the space of possibilities and limitations of dMVPA. In short, neuroscience and related fields need more software tools that match, or exceed, the versatility and ease-of-use of existing traditional MVPA tools. This is the goal of the DeLINEATE toolbox (Deep Learning In Neuroimaging: Exploration, Analysis, Tools, and Education), which we introduce below.

### 2.4 Pros, cons, and caveats of dMVPA

#### Pro: Potentially greater suitability for complex, many-featured datasets

As discussed earlier, one great promise of dMVPA is the potential to unearth more fine-grained patterns in neuroscience data than the simpler (and commonly linear) techniques of traditional MVPA. However, a fundamental principle of statistics is that more powerful (i.e., more complex) models require more parameters^16^, and reliably estimating more parameters requires larger input datasets. Hence, why deep learning and Big Data are commonly associated with each other. Unlike, say, the Google Images team, most neuroscientists are unfortunately not swimming in training data for sophisticated machine learning; neuroscience data are frequently “Big,” but more from features^17^ than from number of examples^18^. Of course, in deep learning (and most statistical analyses), the inverse situation is usually more desirable: A relatively large ratio of examples to features.

Potential solutions to the too-many-features problem include finding ways to intelligently select (*feature selection*) or algorithmically condense^19^ the feature set. However, beyond those options, most traditional MVPA techniques do not many choices for constraining the feature set, and in particular lack any built-in ability to take the structure of the input data into account. This is unfortunate because neuroscience data^20^ tend to be highly structured (temporally and spatially) in ways that could be informative for MVPA^21^. DNNs, on the other hand, have numerous potential architectural configurations that can be optimized to take advantage of known structure in the input data. Most notably, certain types of ANN layers (e.g., convolutional layers) can handle multi-dimensional input data, whereas traditional MVPA’s linear classifiers typically just vectorize multi-dimensional inputs. Thus, dMVPA makes it possible to design customized classifiers that are more suited to a particular shape/dimensionality of input data.

#### Caveat

Having more architectural options for structuring and condensing complex input data also leads to a paradox of choice; how can one possibly decide on the best DNN architecture for a given dataset? Unfortunately, dMVPA is still a young field, and we are still working on establishing good heuristics for network architectures to handle many-featured datasets. Also unfortunately, this is not one of those methodological choices where differences between options can be chalked up to rounding error; the wrong dMVPA architecture may completely fail to perform above chance in situations where a superior architecture classifies the data fairly accurately.

#### Con: Many potential types of analysis architecture; many of these carry an increased danger of overfitting

Most conventional MVPA techniques (SVM, SMLR, etc.) have a relatively small number of hyperparameters^22^ to adjust, and those hyperparameters can often either be left at default values or automatically estimated by the algorithm without serious adverse effects on performance. In contrast, the number of possible hyperparameters to adjust in dMVPA is effectively infinite. These hyperparameters include the number of layers in the network, the number of units in each layer, the type of each layer^23^, and any number of additional layer-type-specific hyperparameters that can be separately specified for each layer. Thus, even choosing a starting point for how to construct a dMVPA model can be daunting for inexperienced researchers (and experienced ones, too). Furthermore, thanks to the No Free Lunch (NFL) theorem(s) (Wolpert & Macready, 1997; Shalev-Shwartz & Ben-David, 2014), we know that no estimation- or optimization-based analysis technique will be optimal for every dataset or problem domain, and therefore it is impossible to know *a priori* whether a given analysis technique will be optimal for a particular problem. Put another way, if we knew in advance that a particular analysis technique *were* optimal for our problem, then that technique would necessarily be exquisitely tailored to the problem – which means we would essentially already know the structure of the data perfectly, which obviates the need to conduct the analysis.

Compounding the problem, there is no real upper limit, other than available computing power, to how complex dMVPA models can be allowed to grow^24^. For the current status quo of neuroscience data, most possible dMVPA models would be far too complex; many would even contain more parameters to estimate than there are data points in the input set! It would be inaccurate to say these models would fit the data poorly; rather, they would fit the training data *too* well. It is not uncommon to see a complex dMVPA model effectively memorize its training data, producing perfect classification of the training dataset but extremely poor generalization to a test dataset – the classic problem of overfitting.

#### Caveat

Much as SVMs provide a fairly robust method for classification across a surprisingly wide range of data types and problem domains (though they are rarely truly optimal due to NFL), there is some hope that such “pretty good, most of the time” dMVPA architectures might exist as well. Again, the field is young, but during development of the DeLINEATE toolbox, we have often found that relatively simple dMVPA models, consisting of just 1–2 convolutional layers and 1–2 dense layers^25^, perform comparably to (or better than) the industry workhorse of SVMs. A bit of customization is often required to fit the size and shape of the input dataset, and it can be useful to test out different variations of dMVPA architecture on one portion of the dataset before applying the best-performing architecture to the remaining held-out data, but a satisfactory architecture is typically not too difficult to find without excessive trial-and-error. We have found that after some experience using dMVPA, one begins to develop fairly good intuitions about what kinds of architecture might be best suited to a specific problem, but it is still far from an exact science.

As the field progresses, we hope that it will converge on more heuristics for designing dMVPA architectures that perform as robustly as SVMs across datasets, while still retaining the flexibility and other advantages of dMVPA. Still, for many practical applications, it is less important to identify an optimal model than it is to determine if the data can be reliably classified above chance (Hebart & Baker, 2018). With properly implemented cross-validation, this can often be achieved by a wide variety of architectures (assuming the data do contain enough meaningful signal for reliable decoding), with the accuracy difference between sets of hyperparameters being only a few percentage points. Conversely, if the input data contain only noise with respect to the classification problem, any sane architecture should perform at chance on the test set. Thus, while some trial and error may be necessary before deciding that data cannot be classified, exhaustive model search is seldom required. When possible, it is often helpful to conduct a traditional MVPA to get a ballpark estimate of how a reasonably well-configured dMVPA should be expected to perform.

#### Pro: Intrinsically multiclass classification

One advantage of dMVPA whose value is likely underestimated is that it is straightforward to design a “true” multiclass classifier, whereas most traditional MVPA methods are intrinsically binary. Thus, in traditional MVPA, multiclass decisions must generally be built from a combination of binary classifiers^26^. While there is nothing methodologically wrong *per se* with building multiclass decisions from binary ones, the implications are slightly different than those of a true multi-way decision, which should be taken into account when interpreting results. Furthermore, in some commonly-used MVPA tools (e.g., PyMVPA), the multiclass decision procedure is not always transparent to the end user, which can be a point of confusion. Conversely, dMVPA classifiers are able to consider all classification options simultaneously; as a consequence, it is also trivially easy to obtain meaningful prediction scores across all classes for each example in the testing set, which can then be used in analyses that go beyond simple winner-take-all accuracy measures.

#### Pro/Con: Performance

Performance, in the sense of speed, can be either an advantage or a disadvantage of dMVPA. Although dMVPA network architectures can vary so widely that it is difficult to generalize, *prima facie* dMVPA should typically run slower than traditional MVPA, because the calculations involved in training a dMVPA network are more complex. However, for larger datasets (in terms of numbers of features and/or examples), the performance of traditional MVPA techniques may scale more poorly than dMVPA. (See “Benchmarks” below and Table 1 for details.) Thus, beyond a certain dataset size, dMVPA may be the only feasible choice. Also, because the network architecture of dMVPA can be adjusted, researchers have more options; e.g., whether to employ a simpler network that may not achieve maximum accuracy but runs quickly, versus a more complex network that runs slower.

**Table 1.** Running times on synthetic benchmark datasets, in minutes. We processed a synthetic benchmark dataset with three models: a convolutional neural network (CNN), Sparse Multinomial Logistic Regression (SMLR), and Support Vector Machines (SVM). Average running time is listed in minutes. A few SVM models never converged in any reasonable amount of time and are represented in the table with the infinity symbol ∞. See text for further details.

| **CNN** | Trials/<br>condition | 100 | 400 | 900 | 1600 | 2500 | 3600 | 4900 | 6400 | 8100 | 10000 |
| --- | --- | --- | --- | --- | --- | --- | --- | --- | --- | --- | --- |
| Features |  |  |  |  |  |  |  |  |  |  |  |
| 200 |  | .207 | .231 | .244 | .250 | .258 | .295 | .365 | .363 | .438 | .475 |
| 400 |  | .216 | .227 | .238 | .240 | .249 | .278 | .297 | .322 | .428 | .520 |
| 800 |  | .213 | .231 | .242 | .248 | .262 | .279 | .305 | .341 | .421 | .437 |
| 1600 |  | .202 | .258 | .270 | .282 | .298 | .337 | .367 | .414 | .499 | .532 |
| 3200 |  | .205 | .322 | .357 | .372 | .376 | .430 | .499 | .564 | .683 | .709 |
| 6400 |  | .195 | .326 | .505 | .542 | .657 | .678 | .724 | .900 | 1.18 | 1.15 |
| 12800 |  | .292 | .503 | .914 | 1.20 | 1.61 | 1.99 | 2.35 | 2.97 | 2.17 | 2.28 |
| 25600 |  | .360 | .920 | 1.32 | 2.49 | 3.71 | 7.41 | 7.73 | 9.43 | 13.5 | 12.1 |
| **SMLR** | Trials/<br>condition | 100 | 400 | 900 | 1600 | 2500 | 3600 | 4900 | 6400 | 8100 | 10000 |
| Features |  |  |  |  |  |  |  |  |  |  |  |
| 200 |  | .024 | .049 | .053 | .086 | .081 | .112 | .127 | .137 | .178 | .219 |
| 400 |  | .047 | .326 | .321 | .382 | .292 | .358 | .447 | .484 | .541 | .676 |
| 800 |  | .087 | .329 | 1.49 | 1.74 | 1.70 | 1.77 | 1.89 | 1.94 | 2.19 | 2.57 |
| 1600 |  | .137 | .563 | 1.70 | 5.40 | 7.78 | 9.09 | 10.1 | 10.3 | 10.8 | 10.8 |
| 3200 |  | .194 | 1.37 | 2.53 | 6.06 | 12.6 | 24.4 | 30.0 | 41.3 | 50.8 | 54.2 |
| 6400 |  | .258 | 2.59 | 5.59 | 9.57 | 16.6 | 29.4 | 46.9 | 70.7 | 112 | 140 |
| 12800 |  | .354 | 4.19 | 13.7 | 20.5 | 28.3 | 42.9 | 64.6 | 91.2 | 128 | 170 |
| 25600 |  | .456 | 5.90 | 23.6 | 52.8 | 63.8 | 76.6 | 104 | 144 | 192 | 253 |
| **SVM** | Trials/<br>condition | 100 | 400 | 900 | 1600 | 2500 | 3600 | 4900 | 6400 | 8100 | 10000 |
| Features |  |  |  |  |  |  |  |  |  |  |  |
| 200 |  | .003 | .103 | .516 | 1.09 | 2.28 | 6.37 | 20.6 | 47.5 | 88.1 | 148 |
| 400 |  | .004 | .067 | 1.25 | 3.71 | 5.67 | 16.7 | 65.1 | 155 | 304 | 502 |
| 800 |  | .008 | .090 | .710 | 5.32 | 10.4 | 32.0 | 159 | 439 | 920 | 1663 |
| 1600 |  | .014 | .216 | .950 | 2.76 | 9.70 | 36.9 | 240 | 855 | 2302 | 4794 |
| 3200 | | .031 | .438 | 2.03 | 5.64 | 12.1 | 22.7 | 109 | 1098 | 3168 | ∞ |
| 6400 | | .068 | .892 | 4.32 | 13.2 | 28.3 | 55.2 | 134 | 392 | 2065 | ∞ |
| 12800 | | .134 | 1.85 | 9.11 | 28.0 | 65.5 | 133 | 357 | 1241 | ∞ | ∞ |
| 25600 | | .263 | 3.72 | 18.5 | 57.1 | 138 | 314 | 1048 | 3699 | ∞ | ∞ |

#### Caveat

As alluded earlier, dMVPA’s computational costs can be somewhat offset by parallelization, which is better supported by deep-learning software tools than most traditional MVPA tools. This is true even if parallelizing across CPUs/cores, but especially true if using the computer’s GPU. Results vary widely depending on dataset size, network architecture, and the specific hardware involved, but users might roughly anticipate anywhere from a 5x–100x or more speedup for running dMVPA on a GPU versus a CPU. On one hand, these benefits make dMVPA a more competitive option, speed-wise. On the other hand, GPU-accelerated dMVPA does require more specialized hardware and more human effort setting up the relevant drivers and software packages. While we have striven in our toolbox and documentation to keep this process as painless as possible, it is still more effort than is required to run non-GPU-accelerated analyses; whether that effort is well-spent will heavily depend on individual users and what tasks they are trying to accomplish.

#### Pro: Flexibility of applications

Although our focus has been on dMVPA, we should note that modern neural networks have an ever-increasing number of uses beyond simple classification. For example, one currently popular strategy is to train a model for categorization within some domain (e.g., the contents of a photograph) and then interrogate the model’s intermediate layers, in an attempt to understand what strategy the model is using (Zeiler & Fergus, 2014). Autoencoder-style architectures allow for, e.g., unsupervised learning of feature structure (Xie et al., 2016), feature-sharpening for degraded inputs (Lore et al., 2017), and principled fusion of multimodal data (Ngiam et al., 2011). Deep networks can also be used to implement classification techniques that are not well-suited to traditional MVPA – for example, “transfer learning,” in which a network is initially trained on one dataset, and then refined by training it further on a different dataset. As another example, we have recently explored using deep networks to create “smarter” similarity/distance metrics tailored to particular datasets/applications, unlike traditional formula-based metrics (e.g., Pearson correlation, Euclidean distance), which do not afford such flexibility (Williams et al., 2020). The DeLINEATE toolbox can, with varying degrees of effort, support many of these advanced applications.

#### Con: Field and dependencies are in active development

While the software tools for traditional MVPA will presumably keep receiving periodic updates, the field overall is fairly mature and not changing particularly rapidly. However, deep learning and dMVPA are newer; as such, the techniques and their underlying software tools are continually being updated. This means that documentation can rapidly go out of date, and incompatibilities can arise easily if developers are not careful. We have aspired to make our own toolbox as robust as possible to the changing software landscape, but it is still worth being aware of. Of course, there are mitigating strategies: Users can find one version that works and refuse to update anything, but this deprives them of future enhancements. Alternately, they can continually update, but this makes it harder to exactly replicate earlier work run with previous software versions. If only Python toolboxes (our DeLINEATE toolbox, and the Keras/PyMVPA backends it relies on) are updated, Python’s “virtual environment” feature can be helpful for maintaining different software setups, each in their own containers. But, if later updates require newer hardware drivers, *and* users wish to maintain backward compatibility with their earlier work, they may wish to do what our lab has done: Purchase several small hard drives for each machine, set up a fresh operating system for each new major driver version, and simply reboot from a different boot drive when one wishes to work with current vs. legacy versions of the software.

### 2.5 A brief introduction to network architecture

In an abstract sense, all feedforward^27^ neural networks may be viewed as a collection of mathematical operations to be applied in sequence to an input of some fixed size, along with rules for updating the parameters of those operations during training. In a classic perceptron, the core operations are multiplication (input data times weight values), summation, and then activation (a thresholding operation, traditionally). In a *multi-layer* perceptron network (Figure 2A), this complete multiplication-summation-activation sequence is repeated, with each layer’s outputs becoming the next layer’s inputs. A typical, slightly simplified mental model for such networks treats those multiplication-summation-activation operations as all occurring within a self-contained unit or node, like in a biological neuron; a number of such units in parallel constitutes a layer of the network, and the main free parameter chosen by the designer of the network is the number of units in each layer. However, unlike a biological neuron, in an ANN this set of operations is not immutable – one might opt to omit activation, invert values after every step, or do any other sort of mathematical transformation, at any step of the sequence. One could also adopt a different mental framework in which every individual operation is a layer of the network, such that each layer of a perceptron network expands into three sequential computational layers: a multiplication layer, a summation layer, and an activation layer. In Keras, the Python framework upon which the DeLINEATE toolbox’s deep-learning functionality rests, it is possible to work with either of these conceptualizations – e.g., there are individual layer types that can perform thresholding/activation, but the activation operation can also be specified as an argument of other layer types, with the understanding that activation is applied last, after that layer’s primary operation.

In lay terms, when sufficiently tortured and beaten into submission, contemporary deep learning frameworks can be mangled into performing virtually any kind of mathematical operation or transformation on the input data. A full discussion of all the possibilities could fill several books, and is thus beyond the introductory scope of this paper. However, there are a few broadly useful kinds of operation/layer that are particularly worth understanding; novices to deep learning should focus on understanding the basic gist of these fundamental tropes before getting lost in the details. Here, they are described briefly in broad categories; Keras has several subtypes of each depending on details of the desired implementation.

#### 2.5.1 Classic

Called “Dense” layers in Keras, these are layers made of perceptrons (Figure 2A). They compute weighted sums and apply an activation function. Varying the number of computational units in such a layer allows one to increase (e.g., consider more potential weightings) or decrease (e.g., prune less informative features) the dimensionality of the data as it passes through the layer. By default, these layers are fully-connected, meaning that all outputs from one layer are used as inputs for each computational unit in the next layer of the network. As noted earlier, a neural network made entirely of dense layers is sometimes called a “multi-layer perceptron” network architecture.

#### 2.5.2 Convolutional

Convolutional layers (Figure 2B) may be conceptualized as collections of filters that are swept across (in mathematical terms, convolved with) their input. When used to process 2-D photographic data, their function is often likened to visual neurons, which take input from a spatially restricted receptive field, extract some feature if present, and pass along the result to the next layer of the visual processing hierarchy. For readers familiar with digital image processing, they are essentially like other kinds of digital filters (e.g., a blur filter, an edge detector), except that convolutional layers can work with any dimensionality of data (not just 2-D images) and their parameters are learned over the course of training, rather than being pre-defined. The combination of filter shape and input data structure will determine what kinds of feature may be selected for and passed along as output. For example, if each example of input data is a 32 × 1000 array of EEG voltages (e.g., 2 seconds of 32-channel data sampled at 500 Hz), a set of 1 × 10 filters would be capable of detecting high-frequency patterns within individual channels (in this example, patterns that fit inside a 20 ms time window), but insensitive to lower-frequency or purely spatial patterns. Conversely, a set of 10 × 1 filters could detect patterns distributed across multiple channels, but only those that occur instantaneously. However, one could instead employ, for example, a set of 8 × 20 filters, which would be capable of detecting patterns spread across up to eight adjacent channels over a 40 ms time window. Choices about data structure are consequently more important for this class of layers than for a multi-layer perceptron; the input examples would contain identical information if flattened from 32 channels × 1000 timepoints to a single 1 × 32,000 vector, but the meaning of a 1 × 10 filter bank’s outputs would be very different.

#### 2.5.3 Recurrent

Recurrent layers (Figure 2C) are named for their property of having their outputs fed back into themselves as inputs. By maintaining an internal state determined by previous inputs, recurrent units develop a form of memory for sequential data. For example, a 1 × 10 vector input to a classic dense unit would be combined to a single value in only two steps – multiplying each element of the vector by its weight and then summing the results. If the same vector were fed into a recurrent unit (typically called a cell), the first element would be handled in isolation, but evaluation of the second element would include the output of the cell’s operation on the first element. The result of this would, in turn, update the unit’s state to influence its response to the third element, and so on until each element of the input is consumed. Recurrent networks are frequently used to process natural language data (both audio and text) and in general are considered good choices for timeseries data. In our own work, we have not observed any significant benefit over convolutional layers when working with human neuroscience data, and have found recurrent-based networks to take longer to train than convolutional-based networks; however, these findings are likely highly dependent on details of the dataset and research question. As alluded earlier, for common types of recurrent cells, the recurrency is handled within the cell as a form of internal “memory” that is not visible to the rest of the network, so network architectures using recurrent layers can still be considered broadly “sequential” or feedforward, and are thus supported by our toolbox.

#### 2.5.4 Supporting

This is a broad category of operations that, for various reasons, are generally thought of as secondary or historically baked-in to more interesting operations. In Keras, this includes activation layers, various purely utilitarian data-reshaping or simple mathematical operations, dropout (an operation in which some percentage of a layer’s units are ignored; thought to mitigate overfitting), etc. Some of these operations (e.g., activation functions) can be specified either as distinct layers or as parameters to a primary layer, whereas others (e.g., a layer that downsamples the output of the previous layer via averaging) can only be specified as distinct layers.

#### 2.5.5 Practical advice

The following is a combination of our experience and advice we have received from other colleagues. We hope it is helpful as a starting point, but readers should not feel overly constrained by it. While the modern leaders in image recognition involve dozens of layers (Szegedy et al., 2016), in our experience the aim of dMVPA can typically be accomplished with much smaller networks. When working with minimally-processed fMRI/EEG/eye-tracking data, we have found that a good starting point often consists of 1–2 convolutional layers followed by 2–3 dense layers; based on preliminary results from that architecture, one could add or remove layers, adjust the layers’ sizes, or tweak other hyperparameters. See Figure 2D for an example. For maximal effectiveness and interpretability, consideration should be given to the match between the shape of the per-example input data and shape of convolutional filters (e.g., should the filters look across EEG channels, or only within? If across, are channels arranged to be spatially adjacent in the data?). *Leaky ReLU* is usually our preferred activation function, and we have often found dropout values of ~0.3 in dense layers to be beneficial. We have found *Stochastic Gradient Descent* (*SGD*) with the momentum parameter (classical or Nesterov) set to something on the order of 0.9 to be a generally successful optimizer, although the *Adam* optimizer (Kingma & Ba, 2014) also performs well in some situations^28^. New users are encouraged to experiment with everything and keep track of the results; soon, you will likely develop your own favorite architectures and hyperparameters. Do not be afraid to experiment broadly; dMVPA has some powerful advantages, but we are also in a more exploratory phase for this kind of research, and designing a sufficiently performant dMVPA architecture can take significant trial-and-error. Of course, the extent to which that exploration might constitute *p*-hacking depends on your research aims; if that is a potential concern, you may want to design your analysis based on an independent dataset (e.g., one of the sample datasets included in our toolbox), or consider a split-half design in which one half of your data is used to explore analysis architectures and the other half is used for confirmatory purposes.

## 3 dMVPA: A toolbox

### 3.1 The DeLINEATE Toolbox

One major purpose of the DeLINEATE toolbox is to enable rapid exploration of model architectures/hyperparameters while maintaining an accurate record of what was done and how it turned out. These are conflicting goals in common practice – a researcher attempting to iterate on an analysis is often tweaking a script or working directly with a command-line interpreter, perhaps in a Notebook type environment (Grus, 2018), and discarding fruitless branches of exploration along the way. Maintaining an accurate record of each tweak and its results during such rapid prototyping is not easy, and can take more time and coding discipline than many of us have.

Our solution to this problem was a processing pipeline in which a single JSON (JavaScript Object Notation) format^29^ job configuration file fully specifies an analysis: the input data, how it will be divided for cross-validation and rescaled, the model architecture to be trained and evaluated, and the outputs to be saved (Figure 3A). The toolbox translates this JSON file into Python code to execute the specified analysis (or analyses), and saves all desired outputs into .tsv (tab-separated values) files with names that include a user-defined prefix linking them to the original JSON file. A copy of that original JSON file can also be saved alongside the other output, so that even if the original is subsequently overwritten during the exploration process, the “output” copy remains a pristine record of what was run to create a particular set of results.

**Figure 3.**
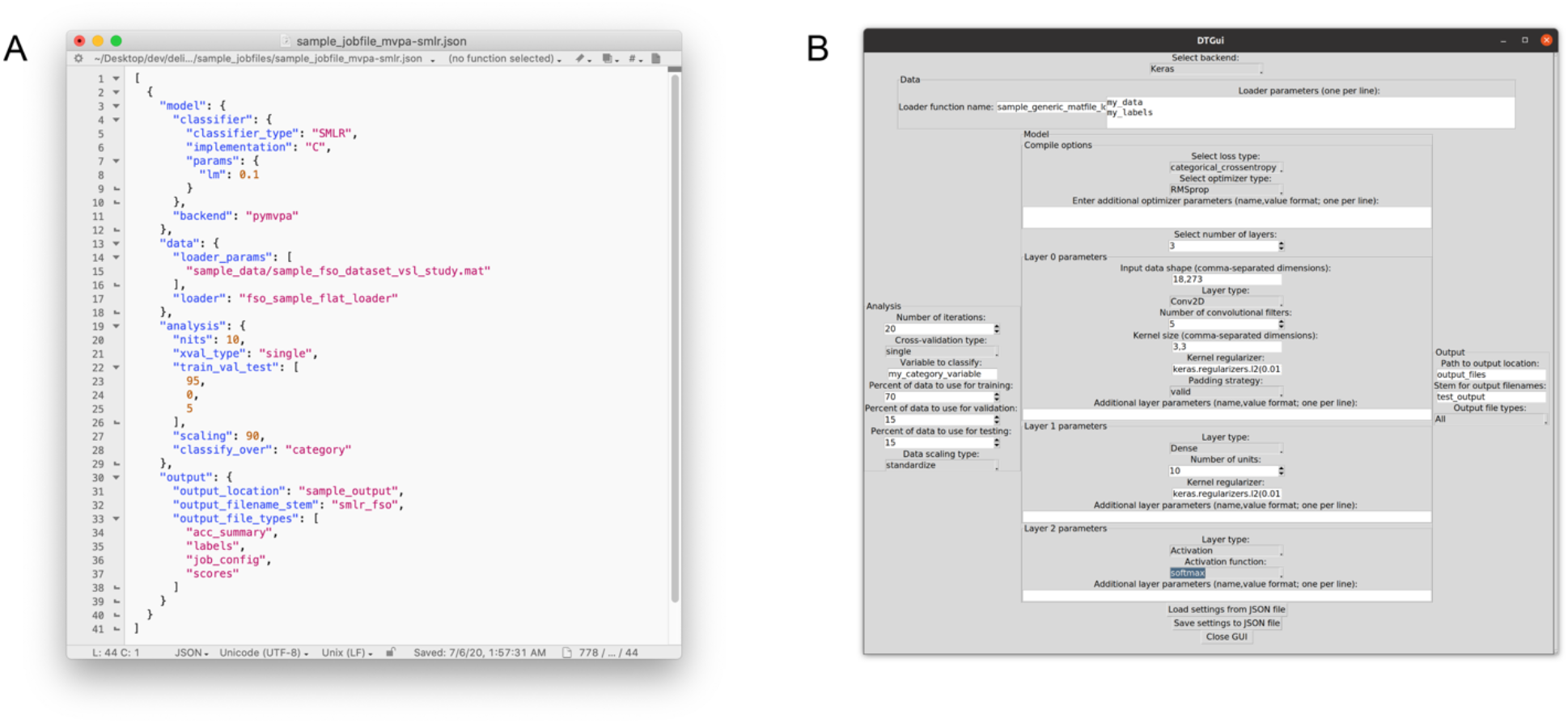
Ways that users can configure an analysis in the DeLINEATE toolbox. (A) Most users will likely configure analyses using a text-based JSON (JavaScript Object Notation) format job file. In this example, the file is open in a generic text-editor program, but JSON-format-specific editing software also exists. Each job has four main sections: “model,” “data,” “analysis,” and “output,” corresponding to the major object types in the toolbox. The file shown is configured to run 10 iterations of a PyMVPA-based SMLR analysis using a sample facescene-object-viewing EEG dataset, using a randomly selected 95% of trials as training data and 5% as test data on each iteration. (B) A basic graphical user interface (GUI) that allows users to configure a job file without having to edit the text directly. The most frequently used options for several common analysis types are available (although editing the text file directly will always allow more flexibility than is possible to express in a GUI). The GUI also contains sections for data, analysis, model, and output, as well as buttons for loading in an existing job file and saving the settings configured in the dialog box to a new JSON file. The settings shown are configured to run 20 iterations of a Keras-based deep learning analysis, using 70% of trials as training data, 15% as validation data, and 15% as test data on each iteration.

A secondary goal was to facilitate comparison of dMVPA approaches to traditional MVPA while, as much as possible, maintaining parity in data handling. To this end, classic MVPA is also supported alongside the dMVPAs that are our primary focus. This is currently implemented with a PyMVPA backend. Traditional MVPA uses the same JSON job file format as dMVPA, as well as similar output file formats, cross-validation/rescaling options, etc., making it a simple task to conduct parallel MVPA and dMVPA on the same data. Currently we support SVM (Support Vector Machine) and SMLR (Sparse Multinomial Logistic Regression) classifiers for traditional MVPA, although our framework is readily extensible to most other classifiers in the PyMVPA toolbox.

For a typical user, the primary entry point to the toolbox is *delineate.py*, a simple script that accepts one or more JSON-format configuration files as arguments, validates their contents, and uses them to create and run one or more analysis job(s). This allows users to run analyses without requiring them to write any code of their own. To further increase accessibility, we have recently developed a simple graphical user interface (GUI) that some find more approachable than a text editor (Figure 3B). GUI users can click on a collection of interactive menus to create properly-formatted job configuration files, which can then be used as input to the main *delineate.py* script. The GUI can also auto-populate selections based on an existing job configuration file for users who have a starting point (such as one of the included sample job files) they wish to modify for future analyses.

For Python-proficient users who want more complex or flexible analysis options, the toolbox can also be used as a Python programming library, and users can write their own code instead of creating JSON files. JSON functionality and code-library functionality can also be mixed-and-matched (e.g., JSON files can be used to create a template analysis, which can then be tweaked and iterated upon with custom code). For users who wish to write their own Python code as well as JSON users who simply want some familiarity with the toolbox’s underlying functionality, we next present a brief overview of the code structure; more detail is available in the toolbox documentation.

### 3.2 DeLINEATE Toolbox structure

The DeLINEATE Toolbox is an object-oriented collection of Python modules, each responsible for a different aspect of the (d)MVPA process. It comprises five main object classes and a small number of supporting files that contain utility functions or facilitate batch analysis. Each main class is housed in a .py file named for that class. In typical usage, the toolbox follows a minimum-import philosophy; to use it as a code library, one simply needs to navigate to its main directory and directly import the desired class file(s). The primary classes are:

1. *DTJob*, responsible for parsing JSON files that define DeLINEATE jobs and passing the appropriate information to constructors for the other object types. In typical usage, a DTJob is responsible for creating one of each other object type and then triggering the DTAnalysis object to actually run the analysis. However, users can also eschew DTJob entirely if they prefer to instantiate the other objects manually in their own Python code.
2. *DTAnalysis*, a parent class that contains one instance each of DTModel, DTData, and DTOutput; it is responsible for coordinating the operations of those other objects. This includes dividing data into training/validation/testing sets, iterating through portions of the data when desired (e.g., to loop through individual subjects), and initiating the model training/testing procedures.
3. *DTModel*, responsible for constructing the model in the appropriate machine learning backend (currently, either Keras or PyMVPA). The “model” in this sense refers either to the artificial neural network (Keras) or an object representing a simpler classifier, e.g., a support vector machine with a linear kernel and parameter C=1 (PyMVPA).
4. *DTData*, responsible for loading the dataset from a data file, storing it, and performing certain operations on it (such as scaling/normalization or slicing it up into smaller training, validation, and/or test subsets).
5. *DTOutput*, responsible for writing analysis results to output files.

The four main sections of a JSON-format job file are the *analysis, model, data*, and *output* sections, which map directly onto the corresponding Python classes; each section contains the parameters necessary to instantiate an object of the appropriate class^30^. Another (purely optional) class, *DTGui*, implements the aforementioned GUI.

### 3.3 Current functionality

#### 3.3.1 Model types and backends

At present, the DeLINEATE toolbox has been used in-house for approximately two years to conduct analyses across a number of studies. It is a high-level toolbox with a flexible, extensible architecture that potentially allows it to sit atop multiple underlying machine-learning libraries. Currently, we support a subset of functionality for two backends: Keras (Chollet et al., 2015) for dMVPA and PyMVPA (Hanke et al., 2009) for traditional MVPA. With our heavy focus on providing a flexible architecture, it is relatively easy to add support for additional backends in the future, as well as enhancing the breadth of support for features of Keras and PyMVPA, enabling new data types to be imported, etc. The relative prioritization of such extensions will be guided by user demand.

#### 3.3.2 Cross-validation

We currently support two approaches to cross-validation. The first is a “universal” approach (specified in configuration files with the name “single”) in which all data are treated as belonging to a single pool, which is randomly divided into training/validation/test sets according to percentages specified in the configuration file. The second divides the data according to some attribute of the samples^31^ and iterates through each value of this property, dividing the data within each iteration into training/validation/test sets (specified in configuration files as “loop_over_sa”). Regardless of which scheme is used, because classification performance can be influenced by a model’s initial conditions^32^, it is common practice to run multiple complete cross-validation iterations in order to ensure a stable estimate of the architecture’s performance. With properly configured input data (see below), these two cross-validation schemes can cover most common MVPA use cases; however, additional schemes can be added in the future according to demand.

#### 3.3.3 Rescaling

Although some MVPA methods are invariant to the scaling of the input data, others, such as many dMVPA applications, require data to be on a certain scale for good classification. The issue is slightly complicated by the need to prevent features of the test data from influencing the training data. We support several methods for rescaling data that avoid this issue by calculating necessary parameters solely on the training data, and using those parameters to adjust validation/test data as well. Again, these methods are readily extensible with additional options, or users can always pre-scale their own data however they like. Currently supported methods are:

1. “percentile”, which identifies the value at a specified percentile of the data and divides all data by that value,
2. “standardize”, which mean-centers and divides all values by the standard deviation of the data,
3. “mean_center”, which subtracts the mean of the data from all values,
4. “map_range”, which translates values into the range between a user-specified minimum and maximum (0 and 1, by default).

#### 3.3.4 Input data and loaders

By far, the most common question we have received from potential users concerns the necessary format for input data. The toolbox operates on, at minimum, one NumPy array and one Python dictionary. The former contains the actual data to be analyzed in a two-or-more-dimensional array, where one dimension represents examples (e.g., trials) and the other dimension(s) are feature dimensions. For instance, an fMRI dataset might be shaped as (examples × voxels), whereas an EEG dataset might be (examples × electrode × timepoint). Higher-dimensional structure is ignored in traditional MVPA and simply collapsed into a 2-D (examples × features) array, as those simple classifiers can only operate on vectors of data. However, dMVPA, when run with an appropriate network architecture, can operate on any dimensionality of data and can potentially take that information into account for classification. If the spatiotemporal structure of the data is meaningful, this may produce superior performance. The Python dictionary contains the metadata needed to interpret the data array, in the form of one or more “sample attributes” (defined earlier; e.g., experimental condition, participant identity) for each sample. These sample attributes may be used as targets for classification (i.e., the class labels to be predicted) or as grouping variables in cross-validation (e.g., for leave-one-subject-out cross-validation).

Data are read into the toolbox by a “loader” Python function specified in the job configuration file. Loaders can reside in a specific subdirectory of the toolbox or in an arbitrary user-specified location. We include several example datasets and corresponding loader functions that should be easily modifiable by researchers to fit their own needs. This is the main place where a typical user *might* need to write their own Python code; because of the many idiosyncratic formats used to store experimental data, some users may need to write a short function to read their files in and reshape them into the expected format. However, if the format is well-supported by NumPy or other Python libraries, these functions can typically be quite short (on the order of 10 lines of code). We also provide generic functions included for data in the NumPy and MATLAB native file formats, which will accept any .mat or .npy file containing one array variable of data examples and at least one variable of sample attributes. Thus, if users are able to save their data in one of those formats beforehand, there may be no need for a custom loader function.

Because neuroscience data vary widely in format, we recognize that a need for additional loader options could still present a barrier to some researchers. We encourage such individuals to reach out to us so that we can offer assistance and expand the range of formats we are able to support natively. On the other hand, the overall flexibility in format means that with just a few lines of code, any dataset that can be represented as a multi-dimensional array is a candidate for analysis with our toolbox, not limited to neuroscience data; for instance, we have used the toolbox to analyze eye-tracking data (Cole et al., under review), photographic images, and more.

#### 3.3.5 Graphical User Interface

As described earlier, the GUI currently allows users to generate a job configuration structure via menu selections and free-entry fields (Figure 3B) that can be auto-populated by loading an existing job file. For frequently used Keras layer types, some reasonable default hyperparameters are provided; however, there are minimal defaults available for less common layer types, and in general it is still recommended for users to have some baseline knowledge of Keras’s workings and hyperparameter options, even when using the GUI. As the number of potential analysis configurations is effectively limitless and this module is a relatively recent addition, error checking is currently somewhat limited. Still, we recognize that a usable GUI is a critical feature for some users, and we expect this to be a primary target for expansion and refinement in upcoming releases.

### 3.4 Availability

All toolbox code is currently hosted at https://bitbucket.org/delineate/delineate and is freely accessible and open-source under the MIT License. There is also a project website at http://www.delineate.it/ that hosts older releases, documentation, links to video tutorials, and more.

### 3.5 Hardware/software requirements

The DeLINEATE toolbox has few software dependencies of its own. However, as noted earlier, it requires either a Keras or PyMVPA backend to perform dMVPA or traditional MVPA, respectively, and those packages have their own corresponding dependencies. Fortunately, both Keras and PyMVPA are well-documented and readily available; we also provide start-to-finish setup guides on the toolbox website. In brief, DeLINEATE is compatible with any recent version of either backend, and in principle can be run on any Python version from 2.7 onward, including all versions of Python 3; however, specific Python version compatibility may depend on which version of Keras/PyMVPA the user is running, and which Python versions *those* libraries are compatible with. The only additional dependency of DeLINEATE is Python support for Tcl/Tk (a graphical interface toolkit) if one wishes to use DTGui; most Python installations include Tcl/Tk libraries, but some might require a separate installation. As Python is available on all major operating systems (Windows, macOS, and Linux), DeLINEATE will also run on any of them, although hardware choices may constrain operating system options.

In terms of hardware, a bare-bones DeLINEATE installation will run on any computer with enough RAM to hold the user’s dataset in memory, as long as the user only wishes to run analyses on the CPU. Traditional MVPA via PyMVPA does not presently employ GPU acceleration, but most dMVPA users will want to enable GPU acceleration for a dramatic increase in speed (see “Benchmarks” below). As Keras relies on the TensorFlow library for its own backend (or the older Theano library; now deprecated in recent Keras versions but still supported by DeLINEATE), which in turn relies on the CUDA (Compute Unified Device Architecture) and cuDNN (CUDA deep neural network) libraries from NVIDIA, effectively this means that an NVIDIA-compatible GPU is required for accelerated dMVPA. Different GPUs will have different compatibility with various versions of CUDA, cuDNN, TensorFlow/Theano, and Keras; however, as long as compatible versions of those tools are installed, DeLINEATE should work with any of them. At the time of writing, we recommend midrange to high-end GPUs from the GeForce 10 series or higher; our lab’s workstations mostly use GeForce GTX 1070 through GeForce GTX 1080 Ti cards, but other users may have higher or lower requirements. Currently, a reasonably powerful workstation for many dMVPA applications could be built from parts for $1500–2000 US^33^, although prices can vary widely depending on users’ specific requirements and budgets. Since no current Apple computers support compatible NVIDIA GPUs, GPU-accelerated dMVPA is currently unavailable on macOS. Generally, for scientific computing, we recommend Linux-based operating systems for their widespread compatibility and open-source nature; however, GPU-accelerated dMVPA will work on Windows as well. In the future, if the macOS/NVIDIA compatibility situation changes, or if DeLINEATE adds support for additional backends, GPU-accelerated dMVPA may become available on macOS.

It has historically been difficult to implement large neural networks without setting up dedicated hardware, largely because the virtualization approaches favored for cloud-based computing do not provide sufficient access to GPUs. However, we have recently seen the emergence of an option that may be useful to those who lack either the budget or the technical confidence to set up their own deep learning environments. Google Colab (https://colab.research.google.com) is a browser-based Python environment akin to Jupyter Notebooks with some access to GPUs. Because the provided environment includes Keras/TensorFlow and allows interaction with files stored on Google Drive, it is relatively straightforward to execute DeLINEATE-based analyses by importing some of the classes and manually calling the method that begins an analysis. An example IPython notebook is provided in the Colab subfolder of the DeLINEATE repository. This approach requires some proficiency in Python and is subject to fluctuating resource limitations, so no promises can be made about speed or stability; however, it may be a good jumping-off point for beginning users wishing to explore the toolbox before investing in their own equipment.

### 3.6 Benchmarks

For both traditional MVPA and dMVPA, performance (both accuracy and computation time) will vary drastically across datasets, hardware, and choice of MVPA classifier or neural network architecture. Thus, the generalizability of any benchmarks is limited. However, to give readers a rough sense of the computational advantages of dMVPA and how running times scale for different dataset sizes, we prepared several datasets and analyzed them with both traditional MVPA and dMVPA. These benchmark datasets emulate the format of an fMRI dataset, but are entirely synthetic. The code to generate them is included in the toolbox.

We simulated datasets with three conditions (classes). Datasets ranged from 200 features (e.g., voxels) to 25,600 features in a doubling progression (200, 400, 800,…). The number of examples (trials) per condition ranged from 100 to 10,000 in the progression: 10^2, 20^2, 30^2,…. Full details are given in the code. Briefly, for each condition, a random signal with the appropriate number of features was generated. Then, supposing for this example that we are generating 900 trials/condition, 30 variations on the “canonical” signal for that condition would be generated by blending the canonical signal with a certain proportion of random noise. Then, for each of those 30 variations, 30 sub-variations were generated by the same process. Although we did not particularly strive for biological verisimilitude, the intent was to somewhat mimic a circumstance where brain patterns had a small number of “true” variations (e.g., if the condition were “faces,” subjects might have slightly different voxel response patterns for different genders/races) as well as trial-to-trial variations due to stimulus exemplar effects and/or measurement noise. To make the classification more challenging, each trial’s signal was also blended with a proportion of the signal of a trial from each of the other two conditions.

The datasets were analyzed with three classifier models: a simple CNN, SMLR, and SVM. The CNN used GPU acceleration (NVIDIA GeForce GTX 1080 Ti), whereas the other models used only the CPU (Intel Xeon X5650 @ 2.67GHz). Each analysis was typically run for 10 iterations (cycles of training/test with different randomly-selected training/test sets) except when running times became prohibitive, in which case the analysis was terminated after as few as five iterations.

Mean running times (Table 1) ranged drastically, from less than one second to several days. As expected, running times for all model types generally increased with greater numbers of features and trials. SVMs had both the shortest and longest running times. Compared to SVMs, SMLR had both a longer shortest running time and a shorter longest running time (i.e., the range was compressed on both ends), and CNNs continued this trend with an even longer shortest running time and a still shorter longest running time (i.e., the range was even more compressed). Notably, the CNN never took less than 10 seconds (largely due to a relatively fixed start-up time for Keras models) but its longest running times, for the most complex datasets, were still under 15 minutes. By comparison, SMLR’s longest running times were over four hours, and SVMs’ were multiple days. (And a few SVM models never converged in any reasonable amount of time.) Thus, as expected, deep learning models were less time-efficient than traditional MVPA for simpler datasets but were vastly more scalable for large datasets.

Benchmark datasets were intended to be classifiable at moderate accuracies but not particularly designed to be benchmarks *of* accuracy, so we do not report comprehensive accuracy results, which could invite misleading extrapolations to real data. However, generally all methods performed above chance, in a comparable range. Typically, the CNN had the lowest accuracy of all three models on datasets with few trials but usually had the highest accuracy with large trial counts, especially when feature counts were low. Conversely, SVM had the highest accuracy when trial counts were low or with very high feature counts, although in those high-feature-count analyses, the SVM running time was long enough to be unusuable in many real-world scenarios. SMLR accuracy almost always fell between CNN and SMLR. Again, we do not expect these accuracies on synthetic data to perfectly reflect performance on real-world data, but they do fit general expectations of how models of varying complexity might be expected to overfit or underfit datasets of varying sizes.

## 4 Discussion

### 4.1 Future development

Toolbox development is ongoing and will largely be steered by community feedback. Current goals include adding support for non-sequential Keras models (e.g., those including feedback connections), transfer learning, model introspection, Generative Adversarial Networks (GANs), and additional built-in data loaders and cross-validation schemes. We also plan to make the GUI more informative and intuitive for users who are less familiar with Keras, and to include some tools for visualization and potentially analysis of results (although this remains an unsettled topic; see Hebart & Baker, 2018, for relevant discussion). Although we have kept discussion in this paper fairly general, information is still liable to go out-of-date quickly due to the rapid pace of deep learning methods development; users are encouraged to consult our website for the most updated details.

### 4.2 Summary

Deep learning continues to grow and offer new possibilities for computation in many areas of research and private industry. While it is being increasingly used in neuroimaging and other neuroscience applications, adoption has been hampered by the complexity of the topic and the lack of approachable software tools. We hope that this tutorial review will help researchers new to deep learning address the former, and that the DeLINEATE software toolbox will help address the latter. In years to come, we expect dMVPA to enable a forward leap in neuroscience discoveries comparable to, or exceeding, that of traditional MVPA over older analyses.

## 6 Nomenclature

ANN: artificial neural network
CNN: convolutional neural network
CUDA^®^: NVIDIA Compute Unified Device Architecture
cuDNN: NVIDIA CUDA^®^ Deep Neural Network library
DeLINEATE: Deep Learning In Neuroimaging: Exploration, Analysis, Tools, and Education
dMVPA: deep multivariate pattern analysis
DNN: deep neural network
GAN: generative adversarial network
JSON: Javascript object notation
ML: machine learning
MVPA: multivariate pattern analysis
SMLR: sparse multinomial logistic regression
SVM: support vector machine

## 7 Conflict of Interest

The authors declare that the research was conducted in the absence of any commercial or financial relationships that could be construed as a potential conflict of interest.

## 8 Author Contributions

KMK, JMW, PCL, and MRJ worked on toolbox code and co-wrote the manuscript. AS and PKR consulted on the analyses and related projects intertwined with toolbox development, and contributed to the writing of the manuscript.

## 9 Funding

This work was supported by NSF Grant CMMI 1719388, Biosensor Data Fusion for Real-time Monitoring of Global Neurophysiological Function awarded to PKR and colleagues, as well as NSF/EPSCoR Grant 1632849, RII Track-2 FEC: Neural networks underlying the integration of knowledge and perception, and NIH P20 GM130461, Rural Drug Addiction Research Center, awarded to MRJ and colleagues. We also received a GPU grant from NVIDIA Corporation. The content is solely the responsibility of the authors and does not necessarily represent the official views of the National Institutes of Health or the University of Nebraska.

## 10 Acknowledgments

We thank Aaron Halvorsen and Hannah Ross for assistance with figure creation and manuscript editing.

## 12 Data Availability Statement

The toolbox code and sample data are available through a Git repository hosted at https://bitbucket.org/delineate/delineate/src/master/. Release versions of the toolbox and additional documentation, as well a link to the Git repository, can be found at http://delineate.it.

## Footnotes

1 Although most of our discussion focuses on fMRI and EEG, as those are the most common techniques in our field of cognitive neuroscience, most points should translate well to related technologies like structural MRI, magnetoencephalography (MEG), or electrocorticography (ECoG), and even to less closely related methods such as extracellular recordings (e.g., from rodents or nonhuman primates).

2 For example, SVMs may take inordinately long to converge on extremely high-dimensional datasets that are handled more easily by deep neural networks. As discussed later, deep networks also have better support for GPU-based parallelization than simpler linear techniques, which can offset their computational costs.

3 The creation of a predictive model that is highly customized to the data used to train the model, but generalizes poorly to new datasets that do not perfectly match the idiosyncrasies of the training data; a significant concern in ML. A good analogy is a bespoke garment perfectly tailored to the contours of a specific individual, which would fit him/her perfectly but look terrible on most others. Conversely, an off-the-rack outfit with a simpler design would fit many individuals of roughly similar proportions reasonably well.

4 Informally defined, the best we might be expected to do in using statistics to explain variance in the data, accounting for the fact that a certain amount of unexplainable variance, aka noise, will always exist.

5 E.g., from better spatiotemporal resolution due to technological improvements; from increasingly large sample sizes, particularly from big-data initiatives such as the Human Connectome Project (Van Essen et al., 2013) and OpenNeuro (formerly OpenfMRI; Poldrack et al., 2013); and simply from the ongoing accumulation of data stockpiles from many years’ worth of research studies.

6 It should be noted up front that the artificial “neurons” used in ML applications bear about as much resemblance to real neurons as a paper airplane bears to a commercial airliner. In both cases, the barest core principles are similar between the pared-down model and the real thing, but little else. However, despite the low resemblance, ANNs can still be extremely useful tools for ML and data processing. Readers are nonetheless cautioned to be as circumspect about over-aggressive comparisons between artificial and real neural networks as they would about buying transatlantic tickets on paper airplanes.

7 As originally conceived, “perceptron” referred to a more complex network of units that could be implemented in a physical machine to produce artificial vision, hence the name. However, the most salient feature that researchers latched onto was the structure of the neural units, and via synecdoche “perceptron” came to be the name of such a unit, so there is some degree of fuzziness around nomenclature and definitions. Here, we use the contemporary sense of “perceptron” to refer to the architecture of an artificial neural unit, rather than the original plan for the physical perceptron machine.

8 The concept of the perceptron is also somewhat looser than the McCulloch-Pitts neuron regarding whether inputs and outputs are constrained to be binary or can be continuously valued, and regarding what kind of thresholding or other function the summed inputs are passed through in order to create the output.

9 Not to mention the ~billion-fold increase in computational power (IBM 704 at 12,000 flops versus a recent desktop GPU at ~11 *tera*flops, for an NVIDIA GeForce GTX 1080 Ti) that helps to make such sophisticated architectures viable.

10 In a biological neural network, one might relate these to a layer of dendrites, a layer of cell bodies, and a layer of axons, but all of those together would comprise a single layer of neurons.

11 Put more formally, single-layer networks cannot solve problems that are not linearly separable, which famously includes the relatively simple XOR function. (For binary inputs A and B, respond “yes” if A is true and B is false, or if A is false and B is true, but respond “no” if A and B have the same value.)

12 Including XOR and many others.

13 As backprop is performed by comparing the performance of a network on a training dataset against an already-known ground truth for that dataset, it is thus considered a form of *supervised learning*, in ML parlance.

14 E.g., the use of ReLU activation functions (Maas, Hannun, and Ng, 2013); new approaches to regularization (Zeiler & Fergus, 2013); and other architectural elements that were available earlier became more prominently used, once sufficient data and computing power existed to use them more effectively (e.g., convolutional network layers).

15 However, alternatives for other GPU architectures do exist, such as the CoreML library used in Apple devices, which use primarily non-NVIDIA GPUs.

16 “Parameters” used in the statistical sense, i.e., numeric values that need to be estimated.

17 In the machine learning sense; for example, the number of voxels in a trial of fMRI data or the number of (electrodes × timepoints) in a trial of EEG data.

18 Also in the machine learning sense, i.e., instances of a set of features that can be assigned a category label. In psychology and neuroscience, such “examples” are generally called “trials” (e.g., of a cognitive task), although in some cases examples may correspond to experimental subjects - an even more limited resource.

19 For example, in techniques like elastic nets (Zou & Hastie, 2005) or SMLR, which use regularization or similar tricks to reduce the number of predictor features.

20 Again, our discussion focuses on neuroscience data, but these techniques, lessons, and software tools can readily be translated to related (or even not-so-related) research fields with similarly-structured datasets and classification problems.

21 For example, it may be useful to condense several spatially adjacent EEG electrodes with similar waveforms into a single data channel. Or, if trying to classify whether a subject is viewing faces or houses, to construct a feature detector that is sensitive to a certain voltage peak (say, the N170; Bentin 1996) but time-invariant within a ~20ms window, to account for trial-to-trial latency variability.

22 This term is less commonly used in the MVPA literature than the ANN literature, but it refers essentially to a parameter of the algorithm set by the user before running the analysis (for example, the amount of regularization), to distinguish those values from plain (non-hyper) parameters, which are the values estimated by the statistical process or model-fitting algorithm.

23 A full rundown of layer types is beyond this article’s scope and better-suited to a general introduction to deep learning, but common types include perceptron-style “dense” layers, “convolutional” layers, “recurrent” layers, and supporting utility layers that calculate simpler mathematical functions; discussed in more detail below.

24 Complexity could be defined many ways, but for now, we will use it mainly to refer to how many parameters (not hyperparameters) need to be estimated for a given model.

25 Technically, these “deep” MVPA networks would not be very deep in terms of how many layers they contain. Still, a fair portion of “deep” learning these days does not use particularly complex network structures; the term now seems to refer more to the contemporary era of ANN-based data analysis than any particular network structure.

26 Typically, if we have classes ABC, the multiclass decision would be made either by training up classifiers “A vs notA,” “B vs not-B,” and “C vs not-C,” or by training up classifiers “A vs B,” “A vs C,” and “B vs C,” and then summing up the scores in favor of each category across classifiers in order to obtain an overall score for that category.

27 “Feedforward” meaning that all outputs from earlier (closer to the input) layers are fed “forward” into later (closer to the output) layers; outputs are never fed back into earlier layers. Feedforward networks are generally easier to work with and design. Our toolbox currently supports only networks with a broadly feedforward design (implemented via the “Sequential” model class in Keras) when using the graphical interface or text-based job files; however, when using it as a collection of Python functions, other network types are possible. One exception is recurrent layers, which feed their output back into themselves; thus networks containing recurrent layers are not strictly feedforward. However, as implemented in our toolbox and the Keras backend we rely on, the recurrency can be viewed as something that recurrent layers handle within themselves; the user does not have to think about this recurrency in terms of their network architecture. From the user’s point of view, the *layers* of the network still follow a feedforward/sequential structure, even if the individual *units* within some layers have recurrency built-in.

28 We realize that all this terminology can be overwhelming at first, but readers unfamiliar with deep learning should try not to feel discouraged by the sheer number of architecture/hyperparameter choices available. Rest assured that it does become more familiar and accessible after some hands-on experience.

29 JSON is a format that allows data structures to be written to plain text files with human-readable syntax. Although not as intuitive as a graphical interface, editing JSON-formatted job files is certainly easier for beginners than writing their own Python code. There are also JSON modules available for many popular text editors and a handful of standalone JSON editing programs to make the task even easier.

30 Although a non-Python-savvy user does not need to know these implementation details, the parity between job file sections and Python classes makes it easy for more experienced coders to switch back and forth between job files and their own Python scripts. As noted above, it is also possible to mix-and-match the two approaches.

31 A “sample attribute,” if you will, which is the terminology used by other MVPA toolboxes for a tag or property associated with each data sample/example. For instance, a subject ID or session ID.

32 Especially for dMVPA; for a given architecture, a classification might sometimes perform well and sometimes at chance depending on the random values assigned to weights at the beginning of training, which is generally a sign that the architecture needs adjusting. Other classification techniques, such as SVMs, are deterministic; as such, they may or may not benefit from multiple cross-validation iterations, depending on dataset and cross-validation scheme.

33 Based on market prices for parts to build a system similar to ours at the time they were built, with an eight-core Intel i7-9700K CPU, GeForce GTX 1070 GPU, 32GB RAM, 1TB SSD primary storage, 4TB HDD secondary storage, and a compatible CPU cooler, motherboard, case, and power supply, for a total of $1750 US. Newer GPUs and other parts have been released since those were built, but pricing for current parts is in a similar range.

## Notes

http://delineate.it

https://bitbucket.org/delineate/delineate/src/master/

### Competing Interest Statement

The authors have declared no competing interest.

